# The role of neural flexibility in cognitive aging

**DOI:** 10.1101/2021.04.22.440855

**Authors:** Eleanna Varangis, Weiwei Qi, Yaakov Stern, Seonjoo Lee

## Abstract

Studies assessing relationships between brain and cognitive changes in healthy aging have shown that a variety of aspects of brain structure and function explain a significant portion of the variability in cognitive outcomes throughout adulthood. Many studies assessing relationships between brain function and cognition have utilized time-averaged, or static functional connectivity methods to explore ways in which brain network organization may contribute to aspects of cognitive aging. However, recent studies in this field have suggested that time-varying, or dynamic measures of functional connectivity, which assess changes in functional connectivity throughout a scan session, may play a stronger role in explaining cognitive outcomes in healthy young adults. Further, both static and dynamic functional connectivity studies suggest that there may be differences in patterns of brain-cognition relationships as a function of whether or not the participant is performing a task during the scan. Thus, the goals of the present study were threefold: (1) assess whether dynamic connectivity (neural flexibility) during both resting as well as task-based scans is related to participant age and cognitive performance in a lifespan aging sample, (2) determine whether neural flexibility moderates relationships between age and cognitive performance, and (3) explore differences in neural flexibility between rest and task. Participants in the study were 423 healthy adults between the ages of 20-80 who provided resting state and/or task-based (Matrix Reasoning) functional magnetic resonance imaging (fMRI) scan data as part of their participation in two ongoing studies of cognitive aging. Neural flexibility measures from both resting and task-based scans reflected the number of times each node changed network assignment, and were averaged both across the whole brain (global neural flexibility) as well as within nine somatosensory/cognitive networks. Results showed that neural flexibility during the task was higher in older adults, and that neural flexibility in Default Mode and Visual networks was negatively related to performance on the Matrix Reasoning task. Resting state neural flexibility was not significantly related to either participant age or cognitive performance. Additionally, no neural flexibility measures that significantly moderated relationships between participant age and cognitive outcomes. Further, neural flexibility differed as a function of scan type, with resting state neural flexibility exhibiting significantly more variability than task-based neural flexibility. Thus, neural flexibility measures computed during a cognitive task may be more strongly related to cognitive performance across the adult lifespan, and are more sensitive to the effects of participant age on brain organization.

## Introduction

Recent work in the cognitive neuroscience of aging has explored the effects of aging on brain network organization and function, both at rest as well as during the performance of cognitive tasks. Generally, when looking at functional connectivity computed across the whole resting state timeseries, studies have found that older adults’ brains tend to be less segregated, less efficiently organized, and have weaker connectivity within networks.^1–3^ Findings linking these age-related differences in network organization at rest to age-related differences in cognitive performance outside the scanner have been relatively limited. They have offered some insight into ways in which resting brain organization/function may explain variability in cognitive performance, but these measure explain a fairly small amount of that variability.^1,3,4^

More recent work in this field has shown that time-varying measures of functional connectivity may account for more, or differential, variability in behavioral outcomes.^5,6^ These time-varying measures typically capture aspects of network reorganization, or differential patterns of interactions between networks, that evolve and change over the course of a resting-state or task-based functional magnetic resonance imaging (fMRI) scan. The most common method for measuring these time-varying patterns is to estimate measures of connectivity during a number of overlapping “sliding windows” in the blood oxygen level-dependent (BOLD) timeseries data. Essentially, network structure/integrity is estimated within each window, with outcome measures reflecting the degree to which these measures change from one window to the next.

While many of these dynamic/time-varying connectivity analyses have focused on younger adults, some studies have extended these analyses to explore aspects of neurocognitive aging. Studies examining time-varying connectivity metrics in older adulthood have found that, at rest, older adults show different patterns of dynamic integration/segregation across multiple brain networks relative to younger and middle-aged adults^7^ and patterns of spontaneous state-switching that may underlie differences in cognitive performance between high- and low-performing older adults^8^. Additionally, one study examining dynamic aspects of functional connectivity during a working memory task found that older adults show reduced ability to reorganize and synchronize functional brain networks to aid in task performance relative to younger adults^9^. However, no studies to date have specifically compared dynamic measures of connectivity across the adult lifespan both at rest as well as during a cognitive task known to be sensitive to age-related changes in fluid cognition. Further, results tying these patterns of dynamic connectivity to cognitive/task performance are limited, providing some evidence that these time-varying connectivity patterns may carry cognitive significance, but with many nuanced relationships between time-varying connectivity and cognitive performance remaining unanswered.

In this vein, a few recent papers explored relationships between time-varying connectivity metrics and behavioral/cognitive outcomes in groups of younger adults and infants. In the first study utilizing a novel method^10^ for detecting dynamic changes in community structure over time in neuroimaging data, the authors collected fMRI data during 3 long motor sequence learning sessions spread out over 5 days^11^ and found that nodes showed flexibility in network membership both within and between experimental sessions, and that flexibility in network assignment during the task predicted later learning. Another study used similar methodology to probe relationships between flexibility observed while performing a working memory task and neuropsychological task performance in younger adults.^12^ The authors found that network flexibility was fairly evenly distributed across most networks during a control task condition, but that during more effortful task conditions, network flexibility was more prominent in frontoparietal and frontotemporal network nodes. Further, the degree of integration among these frontal networks predicted out-of-scanner neuropsychological measures of working memory and executive function, suggesting that the degree of flexibility in these networks during working memory task performance may play a role in explaining variability in performance on similar tasks. Finally, while these studies found promising relationships between task-based neural flexibility and cognitive outcomes in samples of healthy younger adults, a more recent study utilizing the same metric explored similar relationships using infant fMRI scans acquired during natural sleep (“rest”) over the course of the first 2 years of life.^13^ The authors found that, generally, neural flexibility across the whole brain increased over this time period, however neural flexibility remained stable in visual network nodes. Further, they found that neural flexibility in this visual network was negatively associated with cognitive ability at age 5/6. Together, these studies suggest that neural flexibility may provide meaningful information about neural correlates of cognitive function, both when measured during resting or task-based fMRI scans. However, while past studies have examined these patterns in infancy and in younger adults, it is unclear how these measures may be influenced by participant age over the adult lifespan, and whether they may explain age-related differences in cognitive function.

An additional consideration in these studies is whether the relationships between these measures and cognitive/behavioral outcomes differs when neural flexibility is measured at rest versus during a task. Static connectivity studies that assess aspects of network organization and function often find similar effects of age on connectivity when connectivity is assessed when participants are at rest or performing a cognitive task, but these effects may be larger or smaller than those observed at rest depending upon the task being performed. For example, a recent study in our group found that the nature of the in-scanner task had a significant effect on discovery of age effects on connectivity metrics, with these effects appearing to be more evident during tasks of fluid reasoning than during tasks of episodic memory.^14^ Further, in-scanner task performance showed a variety of relationships between fluid reasoning task performance and connectivity within/between networks during these tasks, suggesting that at least some variability in task performance was accounted for by these measures. While this study did not directly compare these patterns to those observed at rest, it shows that task selection can have a strong effect on the presence and magnitude of the effects of age on functional connectivity. Therefore, it is critical to explore whether dynamic functional connectivity measures show similar differences as a function of task performance, and whether these differences play a role in explaining variability in performance on the task as a function of age.

The present study explores measures of neural flexibility in an adult lifespan sample based on both scans collected during rest as well as during a fluid reasoning task, Matrix Reasoning. This task was chosen due to its relatively long scan length as well as its demands on executive function, a domain found in a previous study to show relationships between network-specific measures of neural flexibility and cognitive outcomes. In this manuscript, we evaluated the following hypotheses: (1) global neural flexibility during rest and during an executive function task is associated with age and cognitive abilities; (2) relationships between neural flexibility and cognitive outcomes will be network-specific; (3) neural flexibility will moderate relationships between age and cognitive performance; and (4) differences in neural flexibility measured at rest and during a task will be differentially associated with cognitive performance.

## Materials and Methods

### Participants

The participants were drawn from two ongoing studies at Columbia University Irving Medical Center: the Reference Ability Neural Network (RANN) study and the Cognitive Reserve (CR) study.^15–17^ In the initial telephone screening, participants who met basic inclusion criteria (i.e., right-handed, English speaking, no psychiatric or neurological disorders, and normal or corrected-to-normal vision) were further screened in person with structured medical and neuropsychological evaluations to ensure that they had no neurological or psychiatric conditions, cognitive impairment, or contraindication for MRI scanning. Global cognitive functioning was assessed with the Mattis Dementia Rating Scale^18^ on which a minimum score of 135 was required for retention in the study. In addition, any performance on the cognitive test battery that was indicative of mild cognitive impairment was grounds for exclusion. The studies were approved by the Internal Review Board of the College of Physicians and Surgeons of Columbia University. Additional details about procedures can be found in previous reports.^15,16,19,20^ In the study, 561 participants were enrolled to the study at baseline with at least one reference ability composite score. Among them 423 participants had usable resting (n=403) or matrix reasoning task fMRI (n=292) scans.

### Image Acquisition and Preprocessing

All MR images were acquired on a 3.0T Philips Achieva Magnet. There were two 2-hour MR imaging sessions to accommodate the twelve fMRI tasks as well as the additional imaging modalities. Relevant to the current study, T1-weighted MPRAGE scan was acquired to determine cortical thickness, with a TE/TR of 3/6.5 ms and Flip Angle of 8°, in-plane resolution of 256 ⨯256, field of view of 25.4 × 25.4 cm, and 165–180 slices in axial direction with slice-thickness/gap of 1/0 mm. Resting-state fMRI blood oxygen level-dependent (BOLD) resting state scans were collected with the following parameters: TE/TR: 20/2000 ms; Flip angle: 72°; In-plane resolution: 112 × 112 voxels; Slice thickness/gap: 3/0 mm; Slices: 37. Task-based fMRI BOLD scans were collected with the following parameters: TE/TR: 20/2000 ms; Field of view: 240mm; Flip angle: 72°; In-plane resolution: 112×112 voxels; Slice thickness/gap: 3/0 mm; Slices: 41. Each participant was instructed to lie still with their eyes closed, to not think of anything in particular, and to not fall asleep. Additional structural scans were acquired but not reported in the current study. A neuroradiologist reviewed each subject’s scans; any significant findings were conveyed to the subject’s primary care physician.

The Matrix Reasoning task included in the present analyses required participants to recognize a pattern from a series of pictures and identify the last missing piece of the pattern from among eight options. The task was designed to closely mirror traditional matrix reasoning tasks utilized as part of a standard neuropsychological task battery.^21^ Each trial began with a 24-second fixation cross, followed by the stimulus. If a response was made within the first 11 seconds, the stimulus terminated at exactly 11 seconds; if a response was made after 11 seconds, the stimulus was terminated immediately following the response. If no response was made, the stimulus terminated after 85 seconds and this trial was coded as no response. The minimum number of possible trials was seven, which occurred if the participant required 85 seconds per trial, or if a time-out occurred for every trial. The maximum number of possible trials was 18, which occurred if the participant required 11 or fewer seconds on each trial. There was a 35–second ISI between trials.

Images were preprocessed using an in□house developed native space method.^22^ Briefly, the preprocessing pipeline included slice timing correction and motion correction (MCFLIRT) performed using the FSL package.^23^ All volumes were registered (6 df, 256 bins mutual information, and sinc interpolation) to the middle volume. Frame-wise displacement (FWD)^24^ was calculated from the six motion parameters and root□mean□square difference (RMSD) of the BOLD percentage signal in the consecutive volumes. To be conservative, the RMSD threshold was lowered to 0.3% from the suggested 0.5%. Contaminated volumes were then detected by the criteria FWD□>□0.5□mm or RMSD >0.3% and replaced with new volumes generated by linear interpolation of adjacent volumes. Volume replacement was performed before temporal filtering ^25^. Flsmaths–bptf was used to pass motion□corrected signals through a bandpass filter with cut-off frequencies of 0.01 and 0.09□Hz. Finally, the processed data were residualized by regressing out the FWD, RMSD, left and right hemisphere white matter, and lateral ventricular signals ^26^. Using advanced normalization tools (ANTs), each T1 image was registered to the 2mm MNI template, and the residualized images were warped to the 2mm MNI template.

In addition to the inclusion criteria mentioned above, participants had to have data for at least one cognitive domain (n=523), have provided data for either resting state or Matrix Reasoning fMRI scans (n=462), and had to have no more than 30% motion artifact removal (scrubbing) from resting state and/or task-based fMRI scans (n=423). As a result, n=423 adults had either resting state scans or task-based fMRI (Matrix Reasoning task) after quality checks. In total, there were 403 resting state scans and 292 task scans (Table 1); among them, 272 adults had both rest and task fMRI scans (Supplementary Table 1). The scan length was either 5 (150 volumes) or 9.5 minutes (285 volumes) for resting state fMRI, and 14.3 minutes (430 volumes) for Matrix Reasoning task-based fMRI.

**Table 1.**
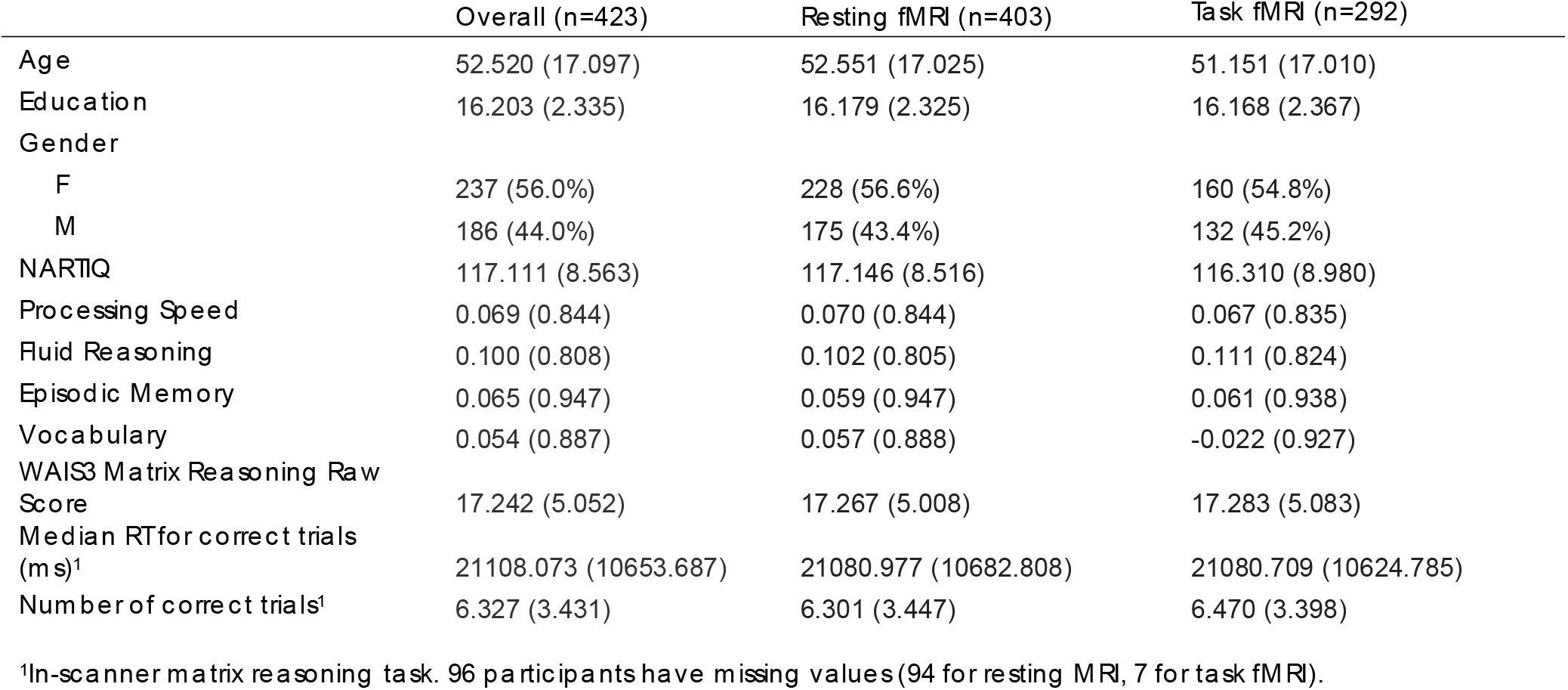
Demographic Characteristics. Means (standard deviations in parentheses) or frequencies (percentage of sample in parentheses) are reported for each sample.

### Neural Flexibility Computation

The nodes were defined using Shen268 ^27^ atlas and the mean time series of each node was extracted. The sliding-window based functional connectivity metrics were computed using Pearson’s correlation coefficients with a window width of 30 volumes and a step size of 1 volume. For dynamic community detection, we employed functions based on generalized Louvain methods with a sliding window approach as described in ^13^. As suggested, we repeated 100 times to get the optimal results. At any given time point each node may have a different community assignment compared to those of the adjacent time points. Given the dynamic community detection results, we defined the neural flexibility (NF) of a node as the number of times that a node changed its community assignment across the sliding windows, normalized by the total number of possible changes. We computed global neural flexibility (GNF) as the average NF over 268 nodes, and network-level NF as the average NF of the nodes based on their membership in 9 Networks (Auditory, Cingulo-opercular, Default-mode, Dorsal-attention, Fronto-parietal, Silence, Sensory-motor, Ventral-attention, Visual) as defined by Power et. al.^28^ and described in Shen et. al.^29^

### Cognitive outcomes

#### Neuropsychological Tests administered out of scanner

Twelve measures were selected from a battery of neuropsychological tests to assess cognitive functioning.^30^ Fluid reasoning was assessed with scores on three different tests: Wechsler Adult Intelligence Scale (WAIS) III Block design task, WAIS III Letter–Number Sequencing test, and WAIS III Matrix Reasoning test. For processing speed, the Digit Symbol subtest from the WAIS-Revised,^31^ Part A of the Trail making test and the Color naming component of the Stroop^32^ test were chosen. Three episodic memory measures were based on sub-scores of the Selective Reminding Task ^33^: the long-term storage sub-score, continuous long-term retrieval, and the number of words recalled on the last trial. Vocabulary was assessed with scores on the vocabulary subtest from the WAIS III, the Wechsler Test of Adult Reading, and the American National Adult Reading Test ^34^. Domain scores were generated by z-scoring performance on each task relative to the full study sample, then average z-scores for tasks within each domain (four domain z-scores: Fluid Reasoning, Processing Speed, Episodic Memory, Vocabulary). Additionally, total number of correct responses on the Matrix Reasoning task was included as an out-of-scanner measure of Matrix Reasoning performance.

#### Computerized tasks administered in the scanner

Twelve tasks from the same 4 domains (Fluid Reasoning, Processing Speed, Episodic Memory, and Vocabulary) were administered in the scanner and their behavioral performance measures were computed using a similar method ^16,19^. In the present analyses, the primary focus will be on performance on the Matrix Reasoning in-scanner task (adapted from Salthouse et. al.^21^). The primary behavioral outcomes for this task are the number of correct trials, and the median correct reaction time.

### Statistical Analysis

For demographic variables, descriptive statistics of mean, standard deviation, frequency and percentage were reported and compared between participants with vs. without neural flexibility measures during rsfMRI sessions, and with and without neural flexibility measures during matrix reasoning task using ANOVA.

We first evaluated the association between age and neural flexibility measures controlling for false discovery rate (FDR) using linear regression. Linear regressions tested the association between NF measures and the four reference ability domain scores (Fluid Reasoning, Processing Speed, Episodic Memory, and Vocabulary) and three matrix reasoning performance measures (out-of-scanner number of correct trials, in-scanner number of correct trials, in-scanner median correct reaction time) adjusting for age, sex and years of education. We further tested the interactions between NF measures and age. For all regression models, standardized regression coefficients, their 95% confidence intervals, and FDR corrected and uncorrected p-values were reported.

In the subsample of n=272 who had both resting and task fMRI data, paired t-tests were performed to examine differences in NF metrics between the two conditions. We tested whether change in neural flexibility between resting and task fMRI is associated with cognitive measures using linear regression, with covariates adjusted.

Lastly, in order to ensure that patterns of results were not exclusively driven by differences in the length of the resting state scan participants completed, all analyses were also performed in just those participants with longer resting state scans (n=291).

Since missing values differ by outcome of interest, data were analyzed using pairwise complete data in all analyses.

### Data availability

MATLAB scripts for neural flexibility computation and additional R markdown files in R 4.0.2. for statistical evaluations and visualizations can be found here: https://seonjoo.github.io/neuralflexibility_brain_submission/

## Results

Details regarding participant eligibility are presented in Figure 1. Demographic characteristics and cognitive performance in the four reference abilities and matrix reasoning tasks for participants included in the present analyses are reported in Table 1. We found that participants who only had resting state data tended to be older 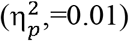, and had higher scores on NARTIQ 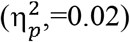 and Vocabulary 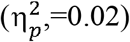 tasks (Supplementary Table 1).

**Figure 1.**
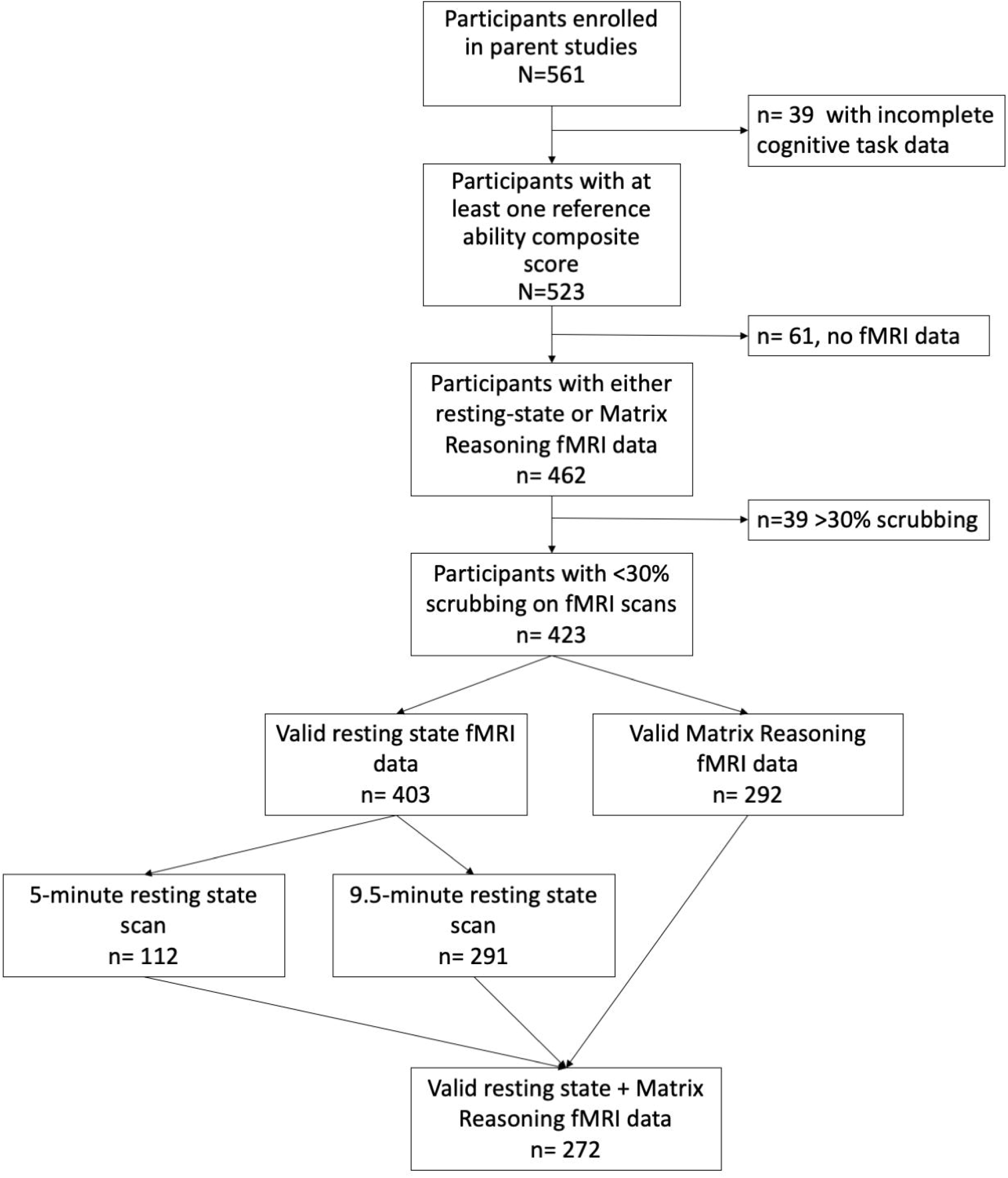
Flow chart of participant enrollment and eligibility.

### Resting State NF

Age was positively correlated with only Auditory network NF; all other correlations between age and NF measures were either nonsignificant, or did not survive multiple comparisons correction (Table 2). Further, resting state NF measures did not significantly predict any cognitive outcomes included in the present study (see Table 3), and did not moderate relationships between age and cognitive performance (see Table 4).

**Table 2.**
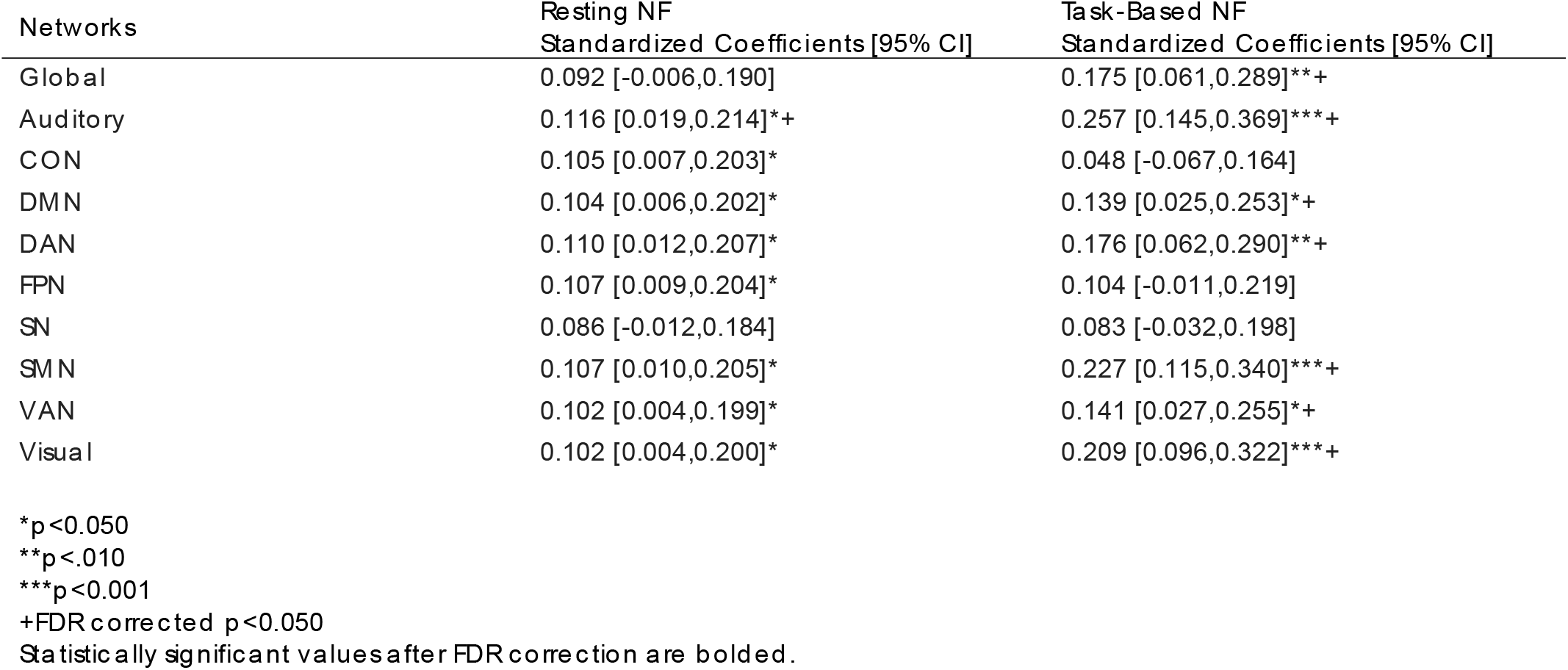
Correlation between NF and Age during resting state and task-based scans.

**Table 3.**
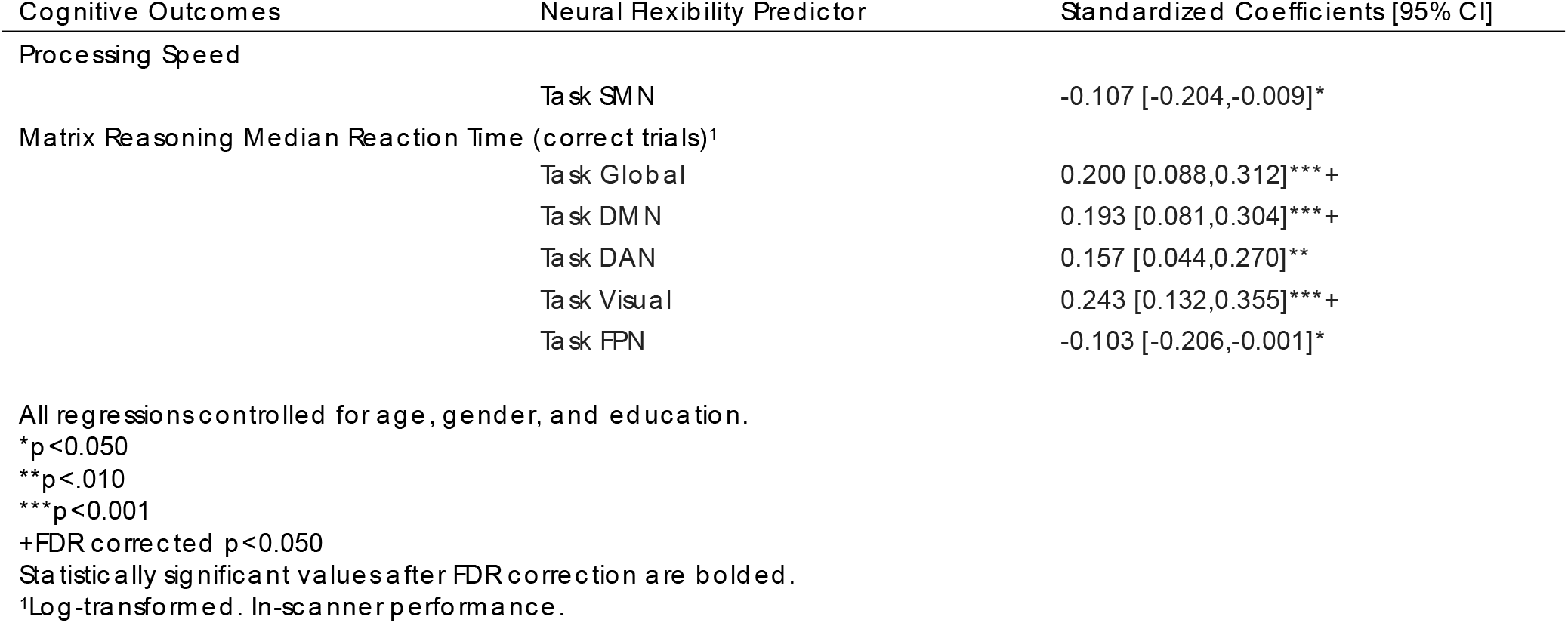
Linear regression between cognitive outcomes and neural flexibility measures. Only significant associations (uncorrected p<0.050) are reported.

**Table 4.** Neural flexibility moderation of cognitive aging. Only significant associations (uncorrected p<0.050) are reported.

| **Cognitive Outcomes** | **Neural Flexibility Predictor** | **Standardized Coefficients [95% CI]** |
| --- | --- | --- |
| **Episodic Memory** | Task Global | -0.120 [-0.225,-0.014]* |
|  | Task AUD | -0.128 [-0.230,-0.026]* |
|  | Task SN | -0.141 [-0.247,-0.036]** |
|  | Task SMN | -0.125 [-0.228,-0.023]* |
| **Matrix Reasoning Median Reaction Time (correct trials)<sup>1</sup>** |  |  |
|  | Task AUD | 0.131 [0.015,0.246]* |
|  | Task CON | 0.127 [0.002,0.252]* |
|  | Task FPN | 0.134 [0.012,0.256]* |

\* $p < 0.050$
\*\* $p < .010$
\*\*\* $p < 0.001$
+FDR corrected $p < 0.050$
Statistically significant values after FDR correction are bolded.
<sup>1</sup>Log-transformed. In-scanner performance.

### Task-Based NF

In task-based NF analyses, age was positively correlated with global NF, as well as network-based NF in the Auditory network, DMN, DAN, DMN, VAN, and Visual network (Table 2). Task-based global neural flexibility was associated with longer median correct reaction time (worse performance) on the matrix reasoning task. (Table 3, β=0.200, 95% CI 0.088 to 0.312, p_fdr_<0.05), controlling for age, gender and education. Network-based associations were found for NF computed based on nodes in the DMN (β=0.193, 95% CI 0.084 to 0.304, p_fdr_<0.05) and the Visual network (β=0.24, 95% CI 0.13 to 0.36, p_fdr_<0.05). Task-based NF measures (global, Auditory, SN, SMN) also moderated age-associated memory decline, such that age-associated memory decline is larger in participants with higher neural flexibility, however these interactions did not survive multiple comparisons correction (Table 4).

### Rest vs. Task

In the subset of participants with both resting and task-based scans (n=272), we compared the neural flexibility measures between rest and task using paired t-tests. The neural flexibility during the task was lower in the DAN and Visual network, and higher in the Salience network than during rest (Table 5; Figure 2). The neural flexibility difference between resting and task was not associated with age, NARTIQ, or task performance (p’s>0.05). Further, neural flexibility in resting scans showed high variability across all networks relative to task-based neural flexibility (Table 5).

**Table 5.** Paired t-tests and variance tests of neural flexibility measures between resting state and task-based scans.

| Neural Flexibility Measure | Mean Within-Subject Difference<br>(Task-Rest) [95% CI] | Cohen's d | Variance Ratio<br>(Rest/Task) [95% CI] |
| --- | --- | --- | --- |
| Global | 0.002 [-0.001,0.004] | 0.084 | 6.750 [5.317,8.569] ***+ |
| Auditory | -0.003 [-0.006,0.000] | -0.118 | 2.456 [1.934,3.117] ***+ |
| CON | 0.002 [-0.001,0.005] | 0.09 | 4.459 [3.512,5.660] ***+ |
| DMN | 0.002 [-0.001,0.005] | 0.082 | 6.503 [5.122,8.255] ***+ |
| DAN | **-0.010 [-0.013,-0.007] \*\*\*+** | **-0.392** | 3.542 [2.790,4.497] ***+ |
| FPN | 0.001 [-0.002,0.003] | 0.025 | 4.984 [3.926,6.327] ***+ |
| SN | **0.004 [0.001,0.007] \*\*+** | **0.181** | 5.775 [4.549,7.332] ***+ |
| SMN | -0.001 [-0.004,0.002] | -0.047 | 3.819 [3.009,4.849] ***+ |
| VAN | 0.002 [-0.001,0.005] | 0.092 | 3.054 [2.406,3.877] ***+ |
| Visual | **-0.014 [-0.017,-0.011] \*\*\*+** | **-0.549** | 3.738 [2.944,4.745] ***+ |

\*p<0.050
\*\*p<.010
\*\*\*p<0.001
+FDR corrected p<0.050
Statistically significant values after FDR correction are bolded.

**Figure 2.**
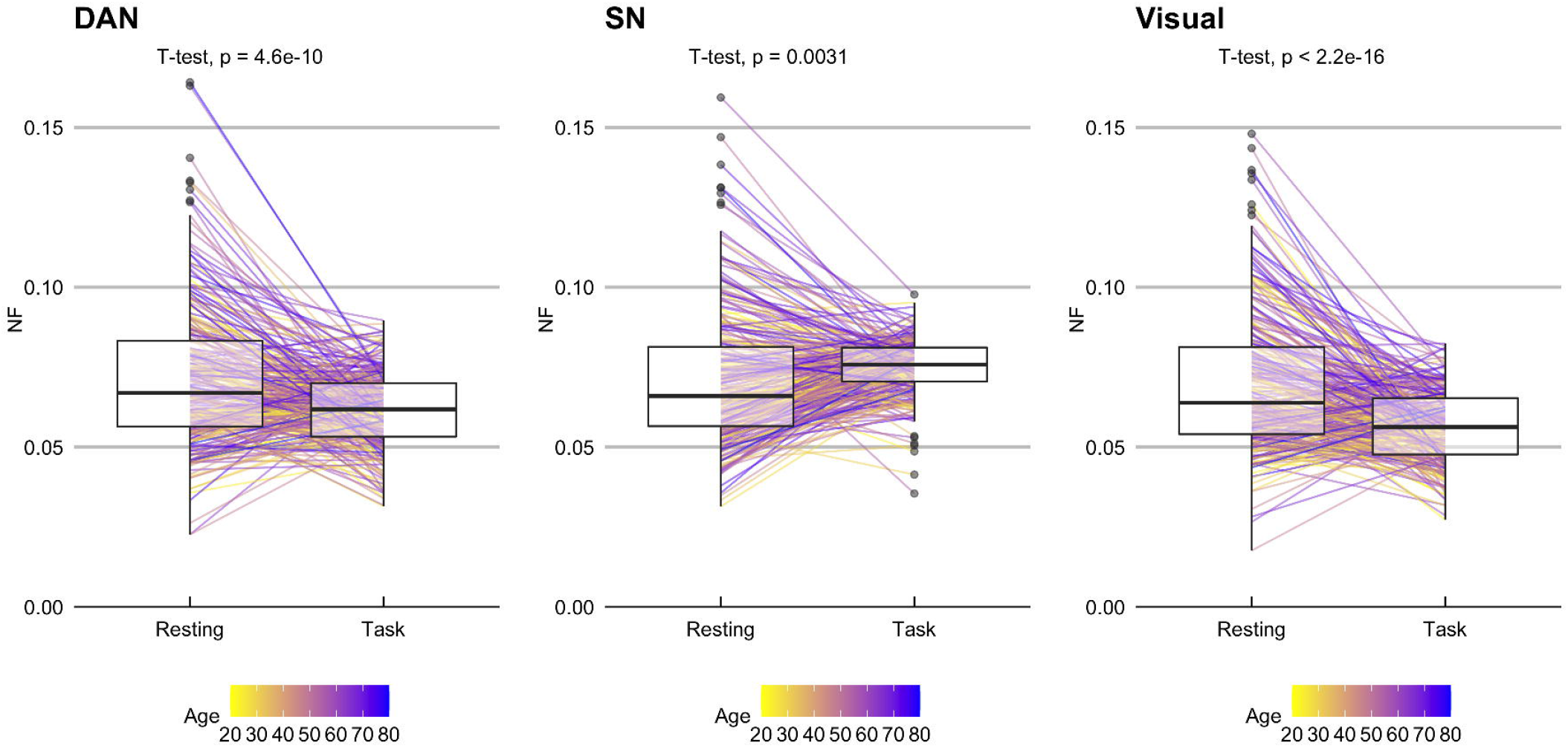
Neural flexibility differences between rest and task.

### Sensitivity Analysis

All analyses reported above were re-calculated in just those individuals with longer resting state scans (n=291). Primary patterns of results described above were largely replicated: (1) age was positively associated with task-based NF, but not resting state NF (Supplementary Table 3); (2) task-based NF in the visual network was positively associated with Matrix Reasoning RT (Supplementary Table 4); (3) NF did not moderate effects of age on cognitive outcomes measures (Supplementary Table 5); and (4) task-based NF differed from resting state NF in several networks (Supplementary Table 6).

## Discussion

In the present study, we found that neural flexibility measured during performance of a task was associated with age and performance of both in- and out of-scanner cognitive tasks, but that there was less of an association between resting state neural flexibility and age or cognitive outcomes. This result may appear to be inconsistent with findings from a study by Yin and colleagues^13^ who reported an association between resting state neural flexibility and cognitive performance, but there are key differences in study design which may account for these differing patterns of results. The Yin et. al. study was conducted in a sample of infants, and showed that neural flexibility during sleep (rest) predicted cognitive outcomes several years later. While sleep is an appropriate resting condition in infants, it may not be exactly analogous to an awake, eyes-open resting state scan in adults, thus patterns of connectivity may understandably show fundamental differences between these two states. Further, infancy is a period of development characterized by rapid neural, cognitive, and motor change, while similar changes on the other end of the developmental spectrum (aging) occur over a much more prolonged period of time. Thus, it is likely that relationships between brain function and cognitive/behavioral outcomes are fundamentally different based on the age of participants in the sample. Therefore, while resting state neural flexibility did not appear to show relationships with age (over the adult lifespan) or cognitive outcomes in the present sample, different patterns may emerge in samples assessing these relationships in different age groups or patient populations.

That being said, other studies in adult samples have found relationships between task-based neural flexibility and cognitive outcomes^11,12^ similar to those identified in the present study. However, the relationships we noted were all in the opposite direction of those previously reported in task-based studies; higher task-based neural flexibility was associated with longer reaction times on the task. This may point to the network specificity of some of the observed effects. For example, Braun and colleagues^12^ specifically focused on the role of frontal neural flexibility, and found positive relationships between task-based neural flexibility in these task-related regions and task performance. In the present study, the network-specific relationships between task-based neural flexibility and task performance were limited to the DMN and Visual networks. In static functional connectivity analyses, the DMN is considered to be a task-unrelated network for tasks of executive function, and studies have found that negative correlations between activity in the DMN and that in task-related networks is associated with better task performance (i.e., Grober et. al.^35^). Further, the prior study referenced above that examined neural flexibility in infants found that resting state visual network neural flexibility was negatively associated with cognitive performance years later.^13^ Thus, while the patterns of relationships observed here suggest that neural flexibility is negatively related to cognition, they also highlight the importance of measuring neural flexibility at the network level in order to investigate the network specificity of these effects. Since these negative relationships between neural flexibility and task performance were observed in networks that may not be associated with the cognitive demands of an executive function task, these relationships represent networks whose integration with other (potentially task-related) networks is associated with poorer performance on the task.

Another important consideration in the interpretation of the results obtained in the present study is that, unlike prior studies focusing primarily on younger adults, the present results reflect patterns of relationships between neural flexibility and cognitive performance across the adult lifespan. Therefore, these observed patterns reflect not only relationships between neural flexibility and cognition in younger adulthood, but relationships that persist throughout the adult lifespan. Thus, while these results may differ slightly from those seen in samples of younger adults, they illuminate important mechanisms that might underlie variability in cognitive outcomes as age-related brain and cognitive changes arise. Further, results from the present study also demonstrated that older adults tend to show higher levels of neural flexibility in several networks during a fluid reasoning task. Specifically, older adults tend to show higher neural flexibility in somatosensory networks (auditory, somatomotor, and visual networks), suggesting that participant age affects measures of neural flexibility, especially those computed during a cognitive task. While these measures did not moderate relationships between age and cognitive task performance, thus are not considered to drive age-related differences in task performance, they may reflect differences in sensory processing as a function of age. Such differences in sensory processing have long been hypothesized to drive age-related differences in cognitive status,^36,37^ and while the present results cannot directly speak to the applications of this theory in these data, they may be suggestive of a neural mechanism by which older adults may compensate for sensory deficits that emerge over the lifespan.

One additional strength of the present study was the ability to directly compare neural flexibility during a task as well as during rest within the same lifespan sample. Neural flexibility during the task was lower in the DAN and visual networks, and higher in the salience network from that during rest. Further, the variance in this measure during the task was much smaller than that during rest, a finding which is in line with previous studies finding higher levels of variability in dynamic connectivity during rest compared to that during a task.^38–40^ While variability in and of itself may be informative when it accounts for variability in relevant behavioral outcomes, the variability observed in the resting state neural flexibility data in the present study was not predictive of cognitive performance, and was not driven by participant age, suggesting that it may be less meaningful than task-based data.

The present study, however was not without limitation. One key challenge faced in this study were the two different resting state scan lengths completed by participants; since resting state scan length can affect functional connectivity metrics, this was of concern in the present analyses. However, follow-up analyses in only longer resting state scans revealed a similar pattern of results (see Supplementary Tables 2-6), suggesting that data from the shorter resting state scans were not likely to be confounding results or increasing variability estimates in the analyses presented here. Further, when exploring group differences between participants with resting state vs. rest+task NF data, several demographic factors differed between these two samples. Given that data for the present study were drawn from two different ongoing studies, this difference is likely driven by the different inclusion criteria for the two studies: the RANN study (contributing resting state + task-based data) included adults between the ages of 20-80; the CR study (only contributing resting state data) included only younger and older adults, with older adults being oversampled in order to better model variability in cognitive outcomes in aging. Further, NARTIQ and Vocabulary scores tend to be higher in older age, thus are likely reflective of this older sample. Therefore, differences in participant age based on task-based scan availability are not driven by systematic exclusion from the task-based scans, but rather based on different inclusion criteria for the two studies whose data are being integrated in the context of the present analyses. Another potential limitation of the present study is in using standardized network assignments for atlas nodes in neural flexibility network-based summary measures, rather than individualized network definitions. While networks were individually estimated as part of the calculation of the neural flexibility estimate, networks used to summarize the data were based on previously described and validated network assignments for this parcellation scheme based on the Power et al.^28^ network assignments.^29^ Thus, while these network summary measures may not necessarily be precisely reflective of each individual participant’s network structure, they are based on widely used network assignments that have been shown to reflect average network structure in healthy adults.

Since the data here represent data from a large, healthy lifespan sample, it is likely that many of the results reported here could generalize to other lifespan samples of healthy adults. Further, since some patterns observed occurred after controlling for participant age, some of the relationships between neural flexibility and cognitive performance may also apply to healthy samples of adults of all ages. That being said, since the present sample is only comprised of healthy, nondemented adults, it is unknown whether the relationships between cognitive performance and neural flexibility would replicate in samples of adults with known cognitive impairment or dementia. Further research is needed in this vein to explore whether these measures can be used to predict cognitive performance in adults with cognitive impairment or dementia.

Based on the data in the present study, neural flexibility measures may quantify some aspect of cognition associated with task performance. However, our results suggest that the neural flexibility measures obtained during rest may be less informative in capturing age-related changes in dynamic brain organization, and may not explain a significant amount of variability in cognitive performance in an adult lifespan sample. During a task, higher neural flexibility (i.e. more frequent changes in network assignment) in specific networks was associated with worse cognitive performance, and older participants tend to have higher neural flexibility, especially in somatosensory networks. These network patterns suggest that flexibility in specific networks may be detrimental to task performance, and that generally older adults show more frequent changes in nodal network assignment.

## Abbreviations

AUD: Auditory Network;
BOLD: Blood Oxygen Level-Dependent;
CON: Cingulo-Opercular Network;
CR: Cognitive Reserve;
DAN: Dorsal Attention Network;
DMN: Default Mode Network;
FDR: False Discovery Rate;
fMRI: functional Magnetic Resonance Imaging;
FPN: Fronto-Parietal Network;
FWD: Framewise Displacement;
GNF: Global Neural Flexibility;
NARTIQ: National Adult Reading Test Intelligence Quotient;
NF: Neural Flexibility;
RANN: Reference Ability Neural Network;
RMSD: Root Mean Square Difference;
SN: Salience Network;
SMN: Somatomotor Network;
VAN: Ventral Attention Network;
WAIS: Wechsler Adult Intelligence Scale

