## Supplementary Table for "The role of neural flexibility in cognitive aging"

Supplementary Materials

### **Section 1: Demographic characteristic comparison by missing status.**

**Supplementary Table1.** Demographic characteristic comparison by missing status.

|  | Rest+Task (N=272) | Resting fMRI  only (N=131) | Task fMRI  only (N=20) | p value |
| --- | --- | --- | --- | --- |
| Age | 51.096 (16.896) | 55.573 (16.959) | 51.900 (18.946) | 0.047 |
| Education | 16.129 (2.355) | 16.282 (2.268) | 16.700 (2.536) | 0.514 |
| Gender |  |  |  | 0.492 |
| F | 151 (55.5%) | 77 (58.8%) | 9 (45.0%) |  |
| M | 121 (44.5%) | 54 (41.2%) | 11 (55.0%) |  |
| NARTIQ | 116.301 (8.947) | 118.941 (7.230) | 116.428 (9.667) | 0.015 |
| Speed Attention | 0.067 (0.835) | 0.074 (0.867) | 0.064 (0.851) | 0.996 |
| Reasoning | 0.115 (0.821) | 0.076 (0.774) | 0.058 (0.879) | 0.875 |
| Memory | 0.052 (0.938) | 0.074 (0.969) | 0.172 (0.950) | 0.856 |
| Vocab | -0.023 (0.931) | 0.223 (0.767) | -0.013 (0.896) | 0.031 |
| WAIS3 Matrix Reasoning Raw Score | 17.322 (5.019) | 17.153 (5.001) | 16.750 (5.999) | 0.862 |

**^1^** In-scanner matrix reasoning task. 92 participants in resting fMRI only group have missing values.

### **Section 2: Sensitivity Analysis including only resting-fMRI data with 9.5-minute scans (n=291).**

**Supplementary Table 2.** Demographic information for the subset of n=291 participants with longer resting fMRI scans.

| Overall (N=291) | |
| --- | --- |
| Age | 53.378 (16.926) |
| Education | 16.254 (2.367) |
| Gender |  |
| F | 165 (56.7%) |
| M | 126 (43.3%) |
| NARTIQ | 117.491 (8.666) |

**Supplementary Table 3.** Correlation between NF and Age during resting state and task-based scans. (n=291)

|  | Resting NF | | Task NF | |
| --- | --- | --- | --- | --- |
| Networks | **Standardized Coefficients [95% CI]** | | **Standardized Coefficients [95% CI]** | |
| Global | 0.118 | [-0.001,0.236] | 0.177 | [0.040,0.315]* |
| Auditory | 0.114 | [-0.005,0.232] | 0.269 | [0.135,0.404]***+ |
| CON | 0.133 | [0.015,0.252]* | 0.025 | [-0.114,0.165] |
| DMN | 0.129 | [0.01,0.247]* | 0.145 | [0.007,0.283]* |
| DAN | 0.139 | [0.021,0.258]* | 0.100 | [-0.039,0.238] |
| FPN | 0.14 | [0.022,0.258]* | 0.053 | [-0.087,0.192] |
| SN | 0.113 | [-0.005,0.232] | 0.075 | [-0.064,0.214] |
| SMN | 0.122 | [0.004,0.241]* | 0.223 | [0.087,0.359]**+ |
| VAN | 0.134 | [0.016,0.252]* | 0.117 | [-0.021,0.256] |
| Visual | 0.138 | [0.02,0.257]* | 0.193 | [0.056,0.330]**+ |

*p<0.05;**p<0.01;***p<0.001, + FDR corrected p<0.05.

**Supplementary Table 4.** Linear regression between cognitive outcomes and neural flexibility measures. Only significant associations (uncorrected p<0.05) are reported. (n=291)

| Cognitive Outcomes | Neural Flexibility | Standardized Coefficients [95% CI] | |
| --- | --- | --- | --- |
| Vocab | Task CON | -0.125 | [-0.244,-0.006]* |
| Matrix Reasoning Median Reaction time (correct trials) ^1^ | Global | 0.191 | [0.062,0.321]** |
|  | Task DMN | 0.195 | [0.067,0.324]** |
|  | Task DAN | 0.149 | [0.020,0.278]* |
|  | Task Visual | 0.270 | [0.143,0.398]***+ |
|  | Task CON | -0.139 | [-0.261,-0.018]* |

All regressions controlled for age, gender and education.

*p<0.05;**p<0.01;***p<0.001, + FDR corrected p<0.05.

^1^Log-transformed. In scanner performance.

**Supplementary Table 5.** Neural flexibility moderation of cognitive aging.

| Cognition | Neural Flexibility | Standardized Coefficients [95% CI] | |
| --- | --- | --- | --- |
| Matrix Reasoning Median Reaction time (correct trials) ^1^ | Task Global | 0.150 | [0.014,0.286]* |
|  | Task CON | 0.159 | [0.016,0.301]* |
|  | Task FPN | 0.149 | [0.011,0.287]* |

^1^Log-transformed. In scanner performance.

**Supplementary Table 6.** Paired t-test of neural flexibility measures between task- and resting state scans.

| Neural Flexibility | Mean Within-subject Difference (task-rest) [95% CI] | | Cohen's d |
| --- | --- | --- | --- |
| Global | 0.000 | [-0.004,0.003] | -0.018 |
| AUD | -0.006 | [-0.010,-0.003] ***+ | -0.247 |
| CON | -0.001 | [-0.004,0.002] | -0.041 |
| DMN | 0.000 | [-0.004,0.003] | -0.016 |
| DAN | -0.013 | [-0.017,-0.010] ***+ | -0.545 |
| FPN | -0.002 | [-0.005,0.001] | -0.095 |
| SN | 0.002 | [-0.001,0.005] | 0.084 |
| SMN | -0.005 | [-0.008,-0.001] *+ | -0.187 |
| VAN | -0.001 | [-0.005,0.002] | -0.052 |
| Visual | -0.016 | [-0.020,-0.013] ***+ | -0.649 |

*p<0.05;**p<0.01;***p<0.001, + FDR corrected p<0.05.
